# Post-translational thioamidation of methyl-coenzyme M reductase, a key enzyme in methanogenic and methanotrophic Archaea

**DOI:** 10.1101/121111

**Authors:** Dipti D. Nayak, Nilkamal Mahanta, Douglas A. Mitchell, William W. Metcalf

## Abstract

The enzyme methyl-coenzyme M reductase (MCR), found in strictly anaerobic methanogenic and methanotrophic archaea, catalyzes a reversible reaction involved in the production and consumption of the potent greenhouse gas methane. The α subunit of this enzyme (McrA) contains several unusual post-translational modifications, including an exceptionally rare thioamidation of glycine. Based on the presumed function of homologous genes involved in the biosynthesis of thioamide-containing natural products, we hypothesized that the archaeal *tfuA* and *ycaO* genes would be responsible for post-translational installation of thioglycine into McrA. Mass spectrometric characterization of McrA in a Δ*ycaO-tfuA* mutant of the methanogenic archaeon *Methanosarcina acetivorans* revealed the presence of glycine, rather than thioglycine, supporting this hypothesis. Physiological characterization of this mutant suggested a new role for the thioglycine modification in enhancing protein stability, as opposed to playing a direct catalytic role. The universal conservation of this modification suggests that MCR arose in a thermophilic ancestor.

## Introduction

Methyl-coenzyme M reductase (MCR) is a unique enzyme found exclusively in anaerobic archaea, where it catalyzes the reversible conversion of methyl-coenzyme M (CoM, 2-methylmercaptoethanesulfonate) and coenzyme B (CoB, 7-thioheptanoylthreoninephosphate) to methane and a CoB-CoM heterodisulfide (1, 2):

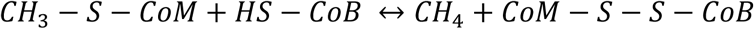

This enzymatic reaction, which is believed to proceed via an unprecedented methyl radical intermediate (3), plays a critical role in the global carbon cycle (4). Thus, in the forward direction MCR catalyzes the formation of methane in methane-producing archaea (methanogens), whereas the enzyme initiates methane consumption in methanotrophic archaea (known as ANMEs for <u>an</u>aerobic oxidation of <u>me</u>thane) in the reverse direction. Together, these processes produce and consume gigatons of methane each year, helping to establish the steady-state atmospheric levels of an important greenhouse gas. MCR displays an α_2_β_2_γ_2_ protein domain architecture and contains two molecules of a nickel-containing porphinoid cofactor, F_430_ (1, 5, 6). The reduced Ni(I) form of F_430_ is essential for catalysis (7), but is highly sensitive to oxidative inactivation, a feature that renders biochemical characterization of MCR especially challenging. As a result, many attributes of this important enzyme remain uncharacterized.

One of the most unusual features of MCR is the presence of several modified amino acids within the active site of the α-subunit. Among these are a group of methylated amino acids, including 3-methylhistidine (or *N*^1^-methylhistidine), S-methylcysteine, 5(S)-methylarginine, and 2(S)-methylglutamine, which are likely installed post-translationally by S-adenosylmethionine-dependent methyltransferases (8, 9). 3-methylhistidine is found in all MCRs examined to date, whereas the presence of the other methylated amino acids is variable among methane-metabolizing archaea (9). A didehydroaspartate modification is also observed in some, but not all, methanogens (10). Lastly, a highly unusual thioglycine modification, in which the peptide amide bond is converted to a thioamide, is present in all methanogens that have been analyzed to date (1, 9).

The function(s) of the modified amino acids in MCR has/have not yet been experimentally addressed; however, several theories have been postulated. The 3-methylhistidine may serve to position the imidazole ring that coordinates CoB. This methylation also alters the p*K_a_* of histidine in a manner that would promote tighter CoB-binding (11). The variable occurrence of the other methylated amino acids suggests that they are unlikely to be directly involved in catalysis. Rather, it has been hypothesized that they tune enzyme activity by altering the hydrophobicity and flexibility of the active site pocket (9, 11). A similar argument has been made for the didehydroaspartate residue (10). In contrast, two distinct mechanistic hypotheses implicate the thioglycine residue in catalysis. The first proposes that thioglycine facilitates deprotonation of CoB by reducing the p*K_a_* of the sulfhydryl group (12); the second suggests that thioglycine could serve as an intermediate electron carrier for oxidation of a proposed heterodisulfide anion radical intermediate (13).

Thioamides are rare in biology. While cycasthioamide is plant-derived (14), most other naturally occurring thioamides are of prokaryotic origin. Among these are the ribosomally synthesized and post-translationally modified (RiPP) peptide natural products thioviridamide (15) and methanobactin (16–18), as well closthioamide, which appears to be an unusual non-ribosomal peptide (19, 20). The nucleotide derivatives thiouridine and thioguanine (21), and two additional natural product antibiotics, thiopeptin and Sch 18640 (22, 23), also contain thioamide moieties. Although the mechanism of thioamide installation has yet to be established, the recent identification of the thioviridamide biosynthetic gene cluster provides clues to their origin (24). Two of the proteins encoded by this gene cluster have plausible roles in thioamide synthesis. The first, TvaH, is a member of the YcaO superfamily, while the second, TvaI, is annotated as a “TfuA-like” protein (24). Biochemical analyses of YcaO-family proteins indicate that they catalyze the ATP-dependent cyclodehydration of cysteine, serine, and threonine residues to the corresponding thiazoline, oxazoline, and methyloxazoline (Fig. 1). Many azoline heterocycles undergo dehydrogenation to the corresponding azole, which are prominent moieties in various RiPP natural products classes, such as linear azol(in)e-containing peptides, thiopeptides, and cyanobactins (25–28). The YcaO-dependent synthesis of peptidic azol(in)e heterocycles often requires a partner protein, which typically is the neighboring gene in the biosynthetic cluster (29–31). Based on enzymatic similarity, the TfuA protein encoded adjacent to the YcaO in the thioviridamide pathway may also enhance the thioamidation reaction, perhaps by recruiting a sulfurtransferase protein, as reported in the biosynthesis of thiouridine and thioguanine (32, 33).

**Figure 1:**
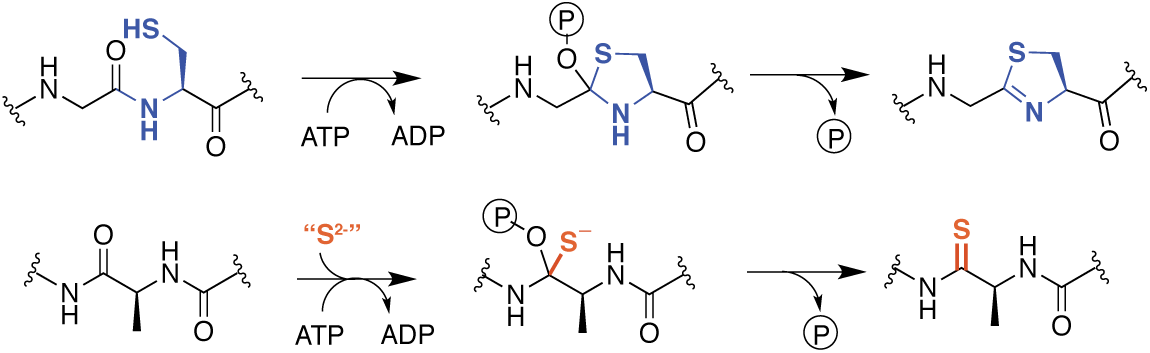
Comparison of reactions catalyzed by YcaO proteins. *Top*, Biochemically characterized YcaO proteins involved in the synthesis of azol(in)e-containing ribosomal natural products catalyze the ATP-dependent cyclodehydration of cysteine, serine, and threonine residues. Shown is the conversion of peptidic cysteine to thiazoline. This reaction proceeds via an O-phosphorylated hemiorthoamide intermediate. *Bottom*, An analogous reaction is believed to occur in the biosynthesis of the thioamide bond in thioviridamide. Rather than an adjacent cysteine acting as the nucleophile, an exogenous source of sulfide (S^2-^) is required for this reaction. TfuA may mediate the delivery of sulfide equivalents for this reaction.

The biosynthesis of the thioglycine in MCR was proposed to occur by a mechanism similar to that used to produce thioamide-containing natural products (9), a prediction made six years prior to the discovery of the thioviridamide biosynthetic gene cluster (24). Given their putative role in thioamidation reactions, it is notable that YcaO homologs were found to be universally present in an early analysis of methanogen genomes, resulting in their designation as “methanogenesis marker protein 1” (34). Genes encoding TfuA homologs are also ubiquitous in methanogens, usually encoded in the same locus as *ycaO*, similar to their arrangement in the thioviridamide gene cluster. Therefore, both biochemical and bioinformatic evidence are consistent with these genes being involved in thioglycine biosynthesis. In this report, we use the genetically tractable methanogen *Methanosarcina acetivorans* to show that installation of the thioamide bond at Gly465 in the α-subunit of MCR requires the *ycaO-tfuA* locus. Significantly, the viability of *ycaO-tfuA* mutants precludes the hypothesis that the thioamide residue is catalytically critical. Instead, our phenotypic analyses support a pivotal role for thioglycine in MCR protein stability.

## Results

### Phylogenetic analyses of TfuA and YcaO in methanogenic and methanotrophic archaea

To examine the possibility that YcaO and TfuA are involved in McrA thioamidation, we reassessed the distribution, diversity, and phylogeny of genes encoding these proteins in sequenced microbial genomes, which today comprise a much larger dataset than when “methanogenesis marker protein 1” was first proposed (34). Significantly, all complete methanogen and ANME genome sequences encode a YcaO homolog, with a few strains encoding two copies (Fig. 2). The YcaO sequences form a distinct, well-supported clade that also includes homologs from ammonia-oxidizing archaea (Fig.2 – Supplemental Figure 1). Excluding the second YcaO in strains that encode two copies, the YcaO tree is congruent with the phylogeny of methanogens reconstructed using housekeeping or *mcr* genes (Fig. 2). Thus, it is likely that YcaO was acquired early in the evolution of methanogens and maintained largely through vertical inheritance, as expected for a trait that coevolved with MCR (35, 36).

**Figure 2:**
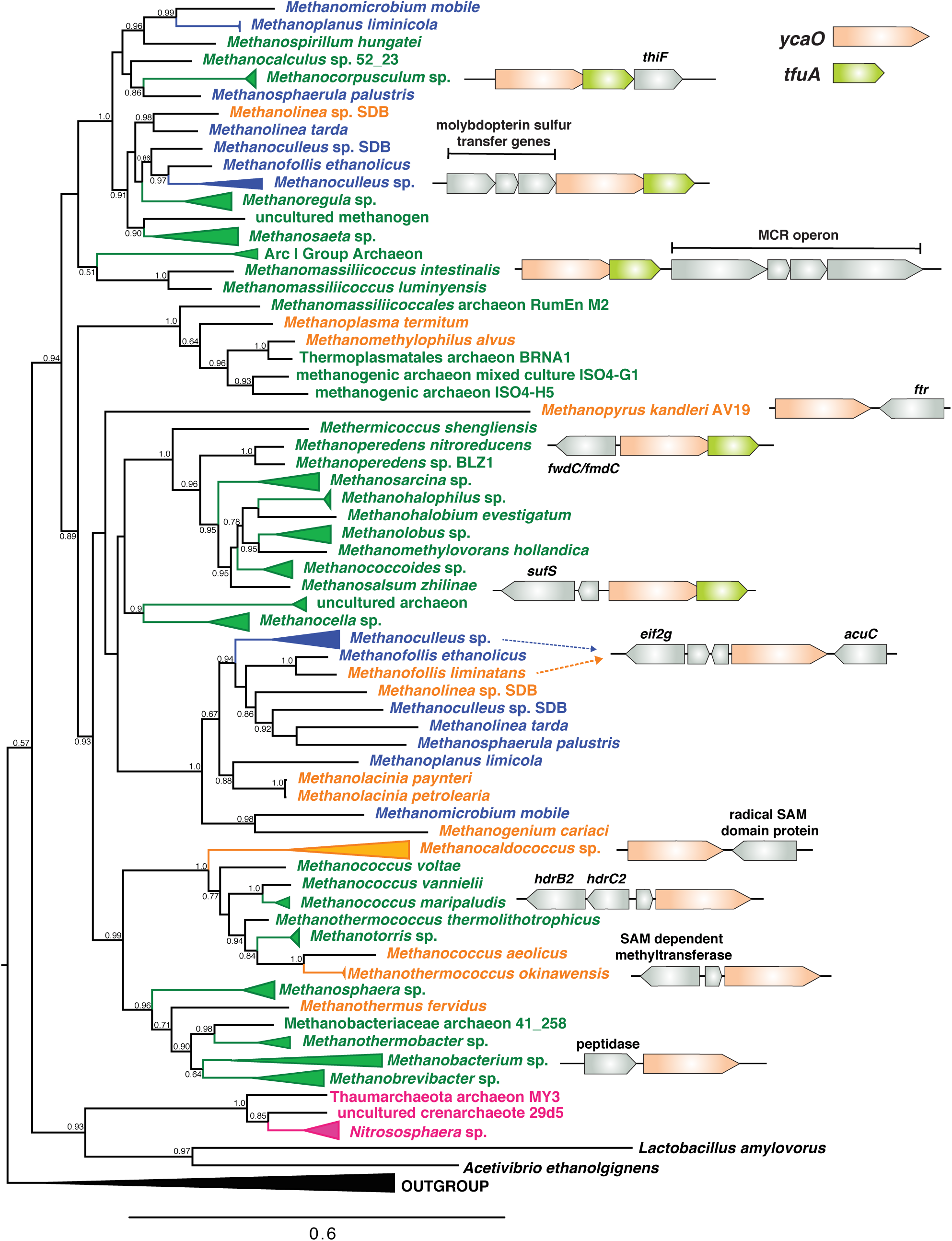
A maximum-likelihood phylogenetic tree of YcaO homologs in archaea. Taxa in green, blue, and orange depict methanogens and <u>an</u>aerobic <u>me</u>thane oxidizing archaea (ANMEs) with sequenced genomes. Taxa shown in green contain a single copy of *ycaO* and *tfuA*, those in blue contain two copies of *ycaO* and one copy of *tfuA*, while those in orange contain one copy of *ycaO*, but do not encode *tfuA*. Taxa shown in pink are other archaea that also encode *ycaO* and *tfuA* homologs. The gene neighborhoods of selected taxa are also depicted, showing the common co-localization with genes annotated as having a role in sulfur metabolism or methanogenesis. The node labels indicate support values calculated using the Shiomdaira-Hasegawa test using 1000 resamples. Support values less than 0.5 have not been shown. The outgroup denotes a variety of bacterial taxa.

TfuA homologs are encoded in the overwhelming majority of MCR-encoding taxa, although a few species lack this gene. The latter include *Methanopyrus kandleri* and *Methanocaldococcus janaschii*, two species previously shown to contain the thioglycine modification (9) (Fig. 2). Thus, TfuA cannot be obligatorily required for thioglycine installation. The archaeal TfuA proteins also fall within a single clade that is largely congruent with the evolutionary history of archaea (Fig.2 – Supplemental Figure 1 and Supplemental Figure 2); however, unlike YcaO, the methanogen clade includes numerous bacterial homologs (Fig.2 – Supplemental Figure 2).

The genomic context of *tfuA* and *ycaO* in methanogens supports a shared or related function, perhaps involving sulfur incorporation and/or methanogenesis (Fig. 2). When both are present, the two genes comprise a single locus in which the stop codon of the upstream *ycaO* gene overlaps with the start codon of the downstream *tfuA* gene, suggesting that they are co-transcribed. In several instances, additional genes involved in sulfur metabolism such as *thiF, sufS*, as well as *moaD, moaE* and *moeB* (involved in molybdopterin biosynthesis) (37, 38) are found in the genomic vicinity. Occasionally, genes encoding enzymes involved in methanogenesis, including the *Methanomassiliicoccus* MCR operon, are found in the neighborhood (Fig. 2).

### TfuA and YcaO are not essential in *Methanosarcina acetivorans*

To test their role in thioglycine installation, we generated a mutant lacking the *ycaO-tfuA* locus in the genetically tractable methanogen *Methanosarcina acetivorans*. Based on the hypothesis that thioglycine may be imperative for MCR activity, and the knowledge that the MCR-encoding operon (*mcrBCDAG*) is essential (39), we first examined the viability of Δ*ycaO-tfuA* mutants using a recently developed Cas9-based assay for gene essentiality (40). This assay compares the number of transformants obtained using a Cas9 gene-editing plasmid with and without a repair template that creates a gene deletion. Essential genes give similar, low numbers of transformants with or without the repair template; whereas non-essential genes give *ca*. 10^3^-fold higher numbers with the repair template. Our data strongly suggest that deletion of the *ycaO-tfuA* locus has no impact on viability (Fig. 3). Several independent mutants were clonally purified and verified by PCR prior to phenotypic characterization (Figure 3 – Supplemental Figure 1).

**Figure 3:**
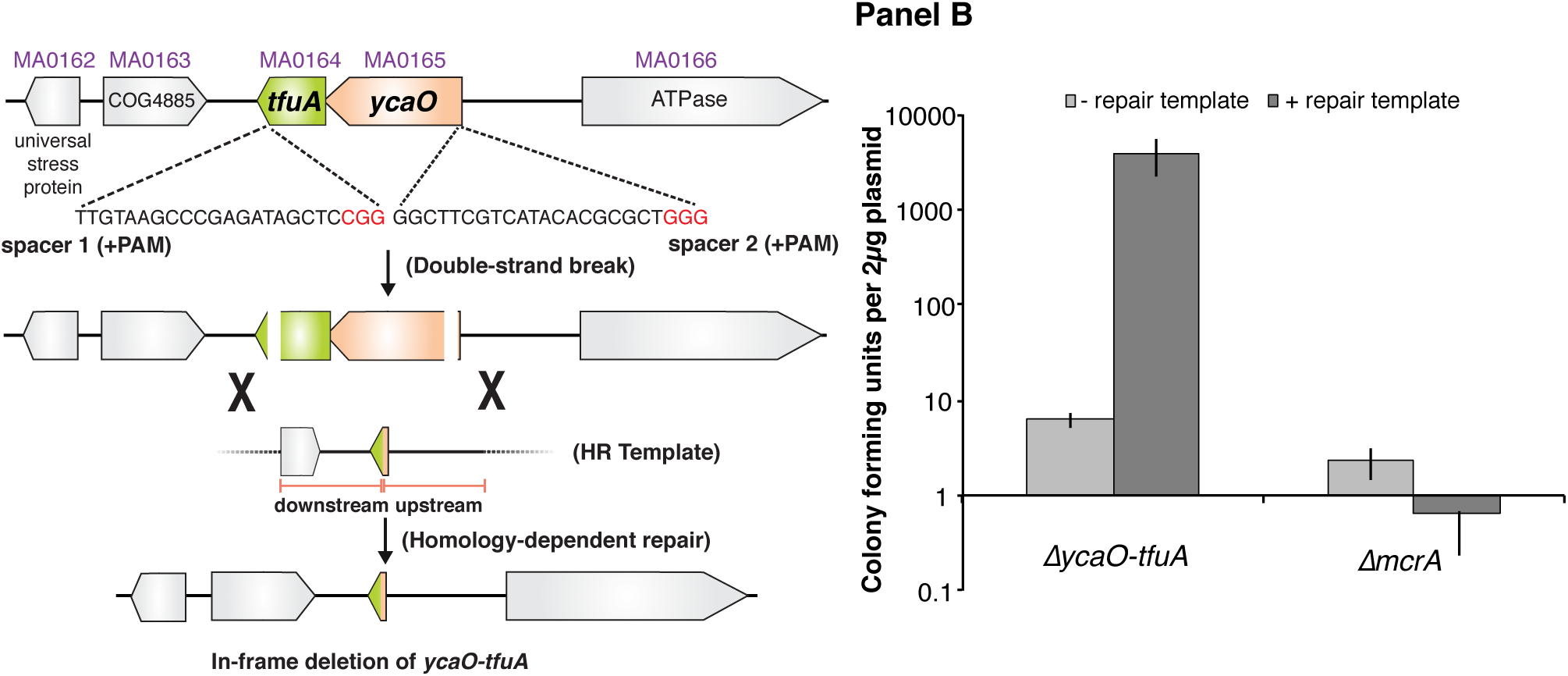
**Panel A**, Experimental strategy for deletion of the *ycaO-tfuA* locus in *M. acetivorans* using a Cas9-mediated genome editing technique. Co-expression the Cas9 protein along with a single guide RNA (sgRNAs) with a 20 bp target sequence flanked by a 3’ NGG protospacer adjacent motif on the chromosome (PAM; in red) generates double-stranded breaks (DSBs) at the *ycaO-tfuA* locus. Repair of this break via homologous recombination with an appropriate repair template (HR Template) generates an in-frame deletion on the host chromosome. **Panel B**, A Cas9-based assay for gene essentiality. Mean transformation efficiencies of plasmids with (dark gray) or without (light gray) an appropriate repair template are shown. The known essential gene *mcrA* is included as a positive control. The error bars represent one standard deviation of three independent transformations.

### The Δ*ycaO-tfuA* mutant lacks the McrA Gly465 thioamide

To test whether YcaO and TfuA are involved in the post-translational modification of Gly465 in McrA, we isolated the McrA protein from cell extracts of the Δ*ycaO-tfuA* mutant and its isogenic parent by excising the appropriate bands from Coomassie-stained SDS-PAGE gels. These proteins were then subjected to in-gel trypsin digestion and the resulting peptides analyzed by matrix-assisted laser desorption-ionization time-of-flight mass spectrometry (MALDI-TOF-MS). The mass spectrum of the parental strain showed a peak at *m/z* 3432 Da (Fig. 4) corresponding to the peptide (_461_LGFFGFDLQDQCGATNVLSYQGDEGLPDELR_491_) with Gly465 being thioamidated (9, 11). The identity of this peptide was confirmed by high-resolution electrospray ionization tandem MS analysis (HR-ESI-MS/MS, [M+3H]^3+^ expt. *m/z* 1144.8608 Da; calc. *m/z* 1144.8546 Da; 5.4 ppm error; Fig. 4 – Supplemental Figure 1A). This peptide also contains the recently reported didehydroaspartate modification at Asp470 (10) and *S*-methylation at Cys472 (8). Consistent with the involvement of TfuA-associated YcaO proteins in thioamide formation, we noted the absence of the *m/z* 3432 Da species in the mass spectrum of a similarly treated Δ*ycaO-tfuA* sample. Instead, a predominant *m/z* 3416 Da species appeared, which was 16 Da lighter, consistent with replacement of sulfur by oxygen. HR-ESI-MS/MS analysis again confirmed the identity of this peptide as being McrA L461-R491 ([M+3H]^3+^ expt. *m/z* 1139.5316 Da; calc. *m/z* 1139.5290 Da; 2.3 ppm error; Fig. 4 – Supplemental Figure 1B). In contrast to the thioglycine-containing wild-type peptide, we observed fragmentation between Gly465 and Phe466 (b_5_ ion, Fig. 4 – Supplemental Figure 1B) in the Δ*ycaO-tfuA* strain. The lack of fragmentation in the wild-type peptide was anticipated based on the greater double bond character of C-N bonds in thioamides (41). The *m/z* 3375 Da species present in both samples corresponds to an unrelated tryptic peptide, McrA Asp392-Arg421 (Fig. 4 – Supplemental Figure 2). We also observed a peak at *m/z* 1496 in both the parent and mutant samples corresponding to a *N*-methylhistidine-containing tryptic peptide (His271-Arg284, Fig. 4 – Supplemental Figure 3A). Thus, thioglycine formation is not a prerequisite for this modification. Additionally, a peak observed *m/z* 1561, corresponding to McrA Phe408-Arg421, shows that Gln420 remains unmodified in *M. acetivorans*, as has previously been observed for *Methanosarcina barkeri* (Fig. 4 – Supplemental Figure 3B) (9).

**Figure 4:**
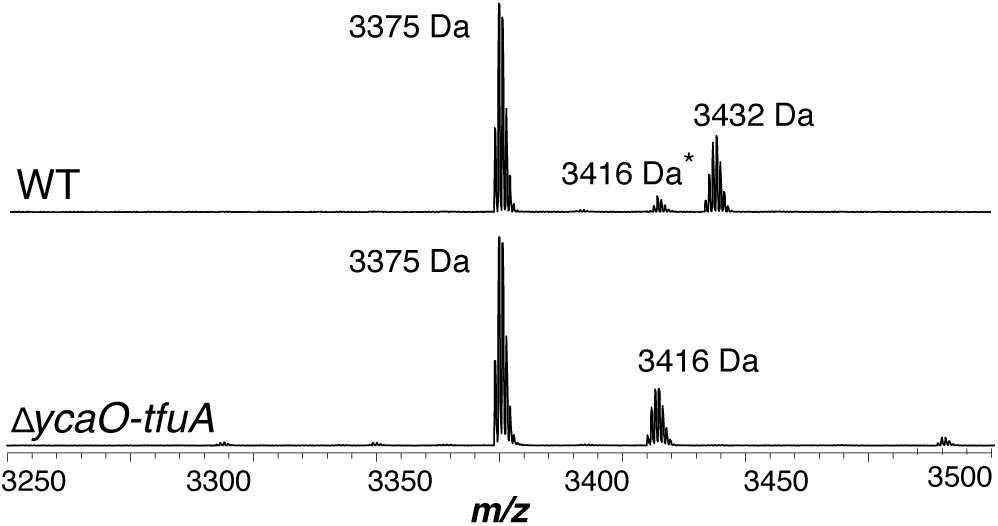
MALDI-TOF MS analysis of MCRα. *Top*, Spectrum obtained from WT tryptic digests containing the thioglycine modified L461-R491 peptide (*m/z* 3432 Da). A small amount of the unmodified L461-R491 peptide (*m/z* 3416 Da) is also observed, which probably arises via non-enzymatic hydrolysis of the thioamide bond as previously reported (9). Bottom, Spectrum obtained from *ΔycaO-tfuA* tryptic digests containing only the unmodified L461-R491 peptide (m/z 3416 Da). The m/z 3375 Da species present in both samples corresponds to MCRα residues D392-R421.

### Growth defects associated with loss of the McrA thioglycine modification

To understand potential phenotypic consequences of losing the thioglycine modification, we quantified the doubling time and growth yield by monitoring changes in optical density during growth of *M. acetivorans* on a variety of substrates (Table 1). While no significant differences were observed during growth on methanol or trimethylamine hydrochloride (TMA) at 36 °C, the Δ*ycaO-tfuA* mutant had substantially longer generation times and lower cell yields on both dimethylsulfide (DMS) and acetate (Table 1). Interestingly, the growth phenotype on methanol medium was strongly temperature-dependent, with no observed differences at 29 and 36 °C, but severe defects for the Δ*ycaO-tfuA* mutant at 39 and 42 °C. Unlike the parental strain, the mutant was incapable of growth at 45 °C (Fig. 5).

**Figure 5:**
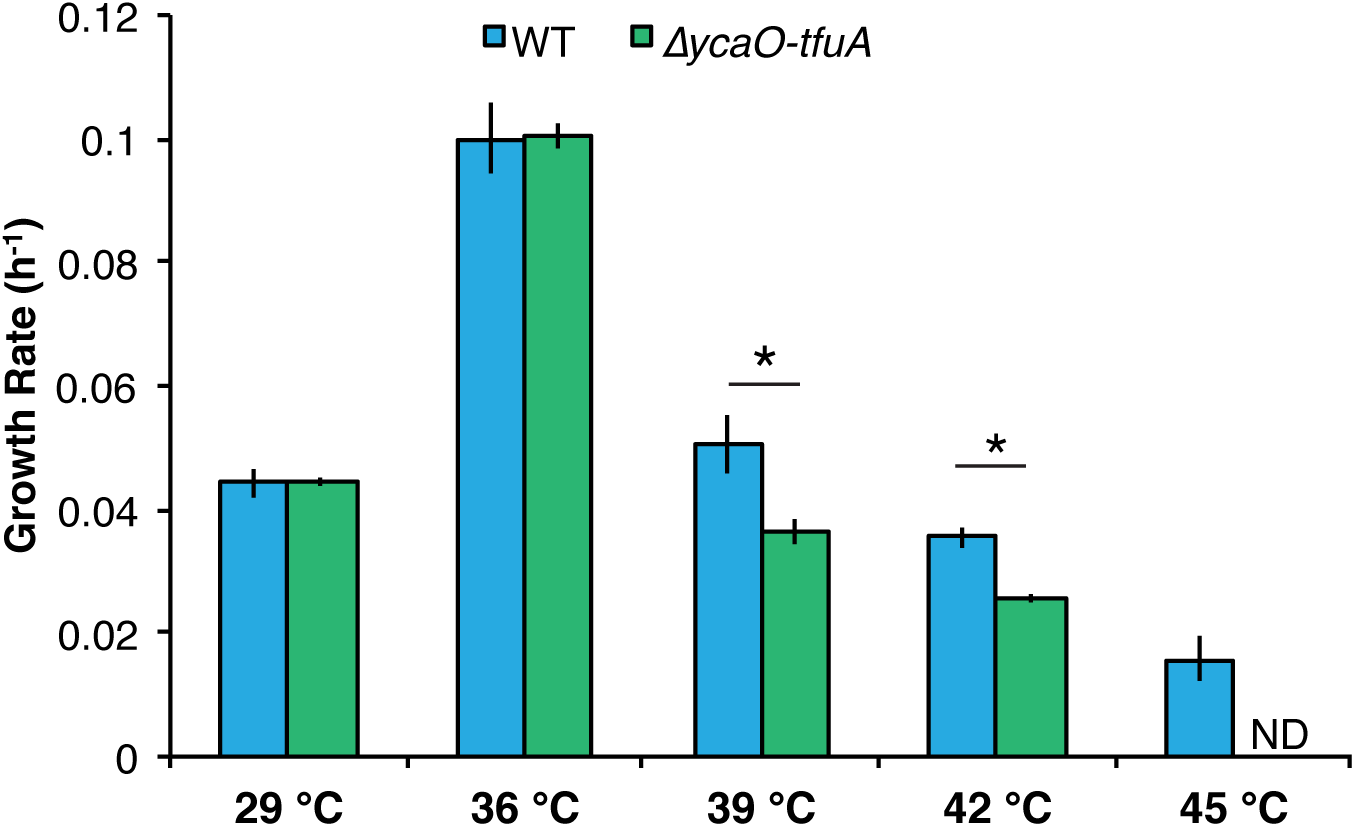
Growth rate of the WWM60 (WT; blue) and the *ΔycaO-tfuA* mutant (green) in HS medium with 125 mM methanol at the indicated temperatures. A statistically significant difference (p < 0.01) using a two-tailed unpaired t-test is indicated with an *.

**Table 1:** Growth phenotypes of WWM60 (WT) and WWM992 (Δ*ycaO-tfuA*) on different substrates at 36°C.

| Substrate<sup>a</sup> | Strain | Doubling Time (h)<sup>b</sup> | p-value | Maximum<br>OD<sub>600</sub><sup>b</sup> | p-value |
| --- | --- | --- | --- | --- | --- |
| **Methanol** | WT | 6.94 ± 0.41 | 0.87 | 5.35 ± 0.31 | 0.26 |
| | $\Delta ycaO$-*tfuA* | 6.89 ± 0.15 | | 5.06 ± 0.39 | |
| **TMA** | WT | 10.92 ± 0.54 | 0.84 | 9.57 ± 0.67 | 0.31 |
| | $\Delta ycaO$-*tfuA* | 10.98 ± 0.43 | | 8.91 ± 0.73 | |
| **Acetate** | WT | 54.23 ± 4.03 | **0.0002** | 0.44 ± 0.01 | **0.0005** |
| | $\Delta ycaO$-*tfuA* | 124.33 ± 12.52 | | 0.35 ± 0.02 | |
| **DMS** | WT | 31.27 ± 1.28 | **<0.0001** | 0.60 ± 0.01 | **<0.0001** |
| | $\Delta ycaO$-*tfuA* | 58.28 ± 1.45 | | 0.44 ± 0.01 | |

<sup>a</sup>TMA, trimethylamine; DMS, dimethylsulfide.
<sup>b</sup>The error represents the 95% confidence interval of the mean for three biological replicates and the p-values were calculated using two-tailed unpaired t-tests.

## Discussion

The loss of the thioglycine modification in the Δ*ycaO-tfuA* mutant suggests that these genes are directly involved in the post-translational modification of McrA. Mechanistically, this conclusion is compatible with biochemical analyses of YcaO homologs. YcaO enzymes that carry out the ATP-dependent cyclodehydration of beta-nucleophile-containing amino acids have been extensively investigated (28). Such cyclodehydratases coordinate the nucleophilic attack of the cysteine, serine, and threonine side chain on the preceding amide carbonyl carbon in a fashion reminiscent of intein splicing (42) (Fig. 1). The enzyme then *O*-phosphorylates the resulting oxyanion and subsequently *N*-deprotonates the hemiorthoamide, yielding an azoline heterocycle. An analogous reaction can be drawn for the YcaO-dependent formation of peptidic thioamides. The only difference is that an exogenous equivalent of sulfide is required for the thioamidation reaction, rather than an adjacent beta-nucleophile-containing amino acid for azoline formation (Fig. 1).

Most YcaO cyclodehydratases require a partner protein for catalytic activity. The earliest investigated YcaO partner proteins are homologs of the ThiF/MoeB family, which are related to E1 ubiquitin-activating enzymes (26, 43). These YcaO partner proteins, as well as the more recently characterized “ocin-ThiF” variety (30), contain a ∼90-residue domain referred to as the RiPP Recognition Element (RRE), which facilitates substrate recognition by interacting with the leader peptide. Considering these traits of azoline-forming YcaOs, it is possible that thioamide-forming YcaOs require the TfuA partner to facilitate binding to the peptidic substrate. Alternatively, TfuA may recruit and deliver sulfide equivalents by a direct or indirect mechanism. In this regard, it is noteworthy that the *ycaO-tfuA* locus can be found adjacent to genes involved in sulfur and molybdoterin metabolism (Fig. 2). Many of these genes encode proteins with rhodanese-like homology domains, which are well-established sulfurtransferases. These enzymes typically carry sulfur in the form of a cysteine persulfide, a non-toxic but reactive equivalent of H_2_S (33, 44). Akin to rare cases of azoline-forming, partner-independent YcaOs, certain methanogens lack a bioinformatically identifiable TfuA (*e.g. Methanopyrus kandleri* and *Methanocaldococcus* sp.). Whether these proteins use an as yet unidentified protein or are truly independent sulfur insertion enzymes remains to be seen. Clearly, further *in vitro* experimentation will be required to delineate the precise role of TfuA in the thioamidation reaction.

The viability of the Δ*ycaO-tfuA* mutant raises significant questions as to the role of thioglycine in the native MCR enzyme, especially considering its universal presence in all MCRs examined to date. We considered three hypotheses to explain this result. First, as we examined the previously suggested possibility that thioglycine modification is involved in enzyme catalysis (9, 13), which would be manifest in a slower reaction rate. As MCR is thought to be the rate-limiting step of methanogenesis (2, 3), this should be reflected in a corresponding decrease in the growth rate on various substrates, with more pronounced defects on substrates that lead to the fastest growth. Significantly, we observed the exact opposite, with the most pronounced defects being observed substrates that support the slowest growth rates in wild-type cells. Thus, loss of the thioglycine modification does not affect the rate-limiting step in growth and therefore is unlikely to result in a slower enzyme. Next, we considered the possibility that thioglycine influences substrate affinity. If so, we expected that differences in metabolite pools caused by growth on different substrates might be reflected in altered growth patterns for the mutant. In particular, C_1_ units enter methanogenesis at the level of *N*^5^-methyl-tetrahydrosarcinapterin (CH_3_-H_4_SPt) during growth on acetate, but at the level of methyl-CoM (45, 46) during growth on DMS, methanol and TMA (Fig. 5 – Supplemental Figure 1; (47, 48)). Significantly, these entry points are separated by an energy-dependent step that is coupled to production or consumption of the cross-membrane sodium gradient. As a result, intracellular levels of methyl-CoM, CoM and CoB could possibly be significantly different depending on the entry point into the methanogenic pathway. Because we observed growth defects on substrates that enter at both points (*i.e*. DMS and acetate), we suspect that the growth deficiency phenotype is unlikely to be related to changes in substrate affinity. However, we recognize that metabolite pools are also affected by the energetics of the methanogenic pathway. Indeed, the growth phenotypes were most severe for DMS and acetate, which are the substrates with the lowest available free energy. The Gibbs free energy (ΔG°’) for methanogenesis from acetate and DMS are -36 kJ/mol CH_4_ (49) and -49.2 kJ/mol CH_4_ (50), respectively, which is considerably lower than the -102.5 kJ/mol CH_4_ and -77.6 kJ/mol CH_4_ for methanol and TMA, respectively (51). Therefore, it remains possible that changes in the substrate pools are responsible for the substrate-dependent growth phenotypes. A third explanation considered was that thioglycine increases the thermal stability of MCR. In this model, unstable MCR protein would need to be replaced more often, creating a significant metabolic burden for the mutant. Consistent with our results, this additional burden would be exacerbated on lower energy substrates like DMS and acetate, especially given the fact that MCR comprises *ca*. 10% of the cellular protein (52). Further, one would expect that a protein stability phenotype would be exaggerated at higher temperatures, which we observed during growth on methanol. Thus, multiple lines of evidence support the idea that the growth-associated phenotypes stemming from the deletion of TfuA and YcaO are caused by decreased MCR stability.

The chemical properties of amides relative to those of thioamides are consistent with our thermal stability hypothesis. Although amides tend to be planar, they have a low rotational barrier and are, thus, conformationally flexible. In contrast, thioamides have higher barriers to rotation and also lessened preference for the *s*-cis conformation (41). Moreover, sulfur has a larger van der Waals radius than oxygen resulting in a thioamide bond length that is ∼40% longer than the amide bond (1.71Å versus 1.23Å) (53), which presents additional steric hindrances to backbone flexibility. Finally, the p*K_a_* of thioamides is lower than the oxygen-containing counterpart, making thioglycine a stronger hydrogen bond donor than glycine (54), which again would reduce conformational flexibility. Taken together, it seems reasonable to conclude that the increased flexibility of the unmodified glycine in the Δ*ycaO-tfuA* mutant results in a protein that is prone to denaturation and enhanced rates of cellular degradation. A conclusive test of this hypothesis will require comprehensive biochemical and biophysical characterization of the MCR variant lacking the thioglycine modification, which is beyond the scope of this work.

Finally, the temperature-sensitive phenotype of the Δ*ycaO-tfuA* mutant has potential implications regarding the evolution and ecology of methanogenic archaea. Based on this result, it seems reasonable to speculate that the thioglycine modification would be indispensable for thermophilic methanogens. It is often posited that the ancestor of modern methanogens was a thermophilic organism (55–57). If so, one would expect the thioglycine modification to be present in most methanogenic lineages, being stochastically lost due to genetic drift only in lineages that grow at low temperatures where the modification is not required. In contrast, if methanogenesis evolved in a cooler environment, one might expect the distribution of the modification to be restricted to thermophilic lineages. Thus, the universal presence of the thioglycine modification supports the thermophilic ancestry of methanogenesis. Indeed, the *ycaO-tfuA* locus is conserved even in *Methanococcoides burtonii*, a psychrophilic methanogen isolated from Ace Lake in Antarctica, where the ambient temperature is always below 2 °C (58). It will be interesting to see whether this modification is maintained by other methanogenic and methanotrophic archaea growing in low temperature environments.

## Materials and Methods

### Bioinformatics analyses

The 1,000 closest homologs were extracted from the NCBI non-redundant protein database using the YcaO amino acid sequence (MA0165) or the TfuA amino acid sequence (MA0164) as queries in BLAST-P searches. The amino acid sequences of these proteins were aligned using the MUSCLE plug-in (59) with default parameters in Geneious version R9 (60). Approximate maximum-likelihood trees were generated using FastTree version 2.1.3 SSE3 using the Jones-Taylor-Thornton (JTT) model + CAT approximation with 20 rate categories. Branch support was calculated using the Shimodaira-Hasegawa (SH) test with 1,000 resamples. Trees were displayed using Fig Tree v1.4.3 (http://tree.bio.ed.ac.uk/software/figtree/).

### Strains, media, and growth conditions

All *M. acetivorans* strains were grown in single-cell morphology (61) in bicarbonate-buffered high salt (HS) liquid medium containing 125 mM methanol, 50 mM trimethylamine hydrochloride (TMA), 40 mM sodium acetate, or 20 mM dimethylsufide (DMS). Cultures were grown in sealed tubes with N_2_/CO_2_ (80/20) at 8-10 psi in the headspace. Most substrates were added to the medium prior to autoclaving. DMS was added from an anaerobic stock solution maintained at 4 °C immediately prior to inoculation. Growth rate measurements were conducted with three independent biological replicate cultures acclimated to the energy substrate or temperature as indicated. A 1:10 dilution of a late-exponential phase culture was used as the inoculum for growth rate measurement. Plating on HS medium containing 50 mM TMA solidified with 1.7 % agar was conducted in an anaerobic glove chamber (Coy Laboratory Products, Grass Lake, MI) as described previously in (62). Solid media plates were incubated in an intra-chamber anaerobic incubator maintained at 37 °C with N_2_/CO_2_/H_2_S (79.9/20/0.1) in the headspace as described previously in (63). Puromycin (CalBiochem, San Diego, CA) was added to a final concentration of 2 μg/mL from a sterile, anaerobic stock solution to select for transformants containing the *pac* (puromycin transacetylase) cassette. The purine analog 8-aza-2,6-diaminopurine (8ADP) (R. I. Chemicals, Orange, CA) was added to a final concentration of 20 μg/mL from a sterile, anaerobic stock solution to select against the *hpt* (phosphoribosyltransferase) cassette encoded on pC2A-based plasmids. *E. coli* strains were grown in LB broth at 37 °C with standard antibiotic concentrations. WM4489, a DH10B derivative engineered to control copy-number of oriV-based plasmids (64), was used as the host strain for all plasmids generated in this study (Supplementary File 1). Plasmid copy number was increased dramatically by supplementing the growth medium with sterile rhamnose to a final concentration of 10 mM.

### Plasmids

All plasmids used in this study are listed in Supplementary File 1. Plasmids for Cas9-mediated genome editing were designed as described previously in (40). Standard techniques were used for the isolation and manipulation of plasmid DNA. WM4489 was transformed by electroporation at 1.8 kV using an *E. coli* Gene Pulser (Bio-Rad, Hercules, CA).

All pDN201-derived plasmids were verified by Sanger sequencing at the Roy J. Carver Biotechnology Center, University of Illinois at Urbana-Champaign and all pAMG40 cointegrates were verified by restriction endonuclease analysis.

### *In silico* design of sgRNAs for gene-editing

All target sequences used for Cas9-mediated genome editing in this study are listed in Supplementary File 2. Target sequences were chosen using the CRISPR site finder tool in Geneious version R9 (60). The *M. acetivorans* chromosome and the plasmid pC2A were used to score off-target binding sites.

### Transformation of *M. acetivorans*

All *M. acetivorans* strains used in this study are listed in Supplementary File 3. Liposome-mediated transformation was used for *M. acetivorans* as described previously in (65) using 10 mL of late-exponential phase culture of *M. acetivorans* and 2 *μ*g of plasmid DNA for each transformation.

### In-gel tryptic digest of McrA

Mid-exponential phase cultures of *M. acetivorans* grown in 10 mL HS medium containing 50 mM TMA were harvested by centrifugation at 5,000 RPM (3,000 × *g*) for 15 minutes in a Sorvall RC500 Plus centrifuge maintained at 4 °C (DuPont, Wilmington, DE) using a SS-34 rotor. The cell pellet was resuspended in 1 mL lysis buffer (50 mM NH4HCO3, pH = 8.0) and harvested by centrifugation at 12,000 RPM (17,500 × g) for 30 minutes in a Sorvall RC500 Plus centrifuge maintained at 4 °C (DuPont, Wilmington, DE) using a SS-34 rotor. An equal volume of the supernatant was mixed with 2x Laemmli sample buffer (Bio-Rad, Hercules, CA) containing 5% β-mercaptoethanol, incubated in boiling water for 10 mins, loaded on a 4-20% gradient Mini-Protean TGX denaturing SDS-PAGE gel (Bio-Rad, Hercules, CA) and run at 70 V until the dye-front reached the bottom of the gel. The gel was stained using the Gel Code Blue stain reagent (Thermo Fisher Scientific, Waltham, MA) as per the manufacturer’s instructions. Bands corresponding to McrA (*ca*. 60 kDa) were excised and cut into *ca*. 1 × 1 mm cubes. The gel slices from a single well were destained with 50% acetonitrile, the corresponding protein was reduced with 10 mM DTT, and digested with 1.5 *μ*g sequencing-grade trypsin (Promega, Madison, WI) at 37 °C for 16-20 hours in the presence of 5% (*v/v*) *n-*propanol. The digested peptides were extracted and dried as described previously (66).

### MS analysis of tryptic peptides

MALDI-TOF-MS analysis was performed using a Bruker UltrafleXtreme MALDI TOF-TOF mass spectrometer (Bruker Daltonics, Billerica, MA) in reflector positive mode at the University of Illinois School of Chemical Sciences Mass Spectrometry Laboratory. The samples were desalted using C-18 zip-tips using acetonitrile and water as solvent system and sinapic acid in 70% acetonitirile was used as the matrix. Data analysis was carried out using the Bruker FlexAnalysis software. For HR-ESI MS/MS, samples were dissolved in 35% aq. acetonitrile and 0.1% formic acid. Samples were directly infused using an Advion TriVersa Nanomate 100 into a ThermoFisher Scientific Orbitrap Fusion ESI-MS. The instrument was calibrated weekly, following the manufacturer’s instructions, and tuned daily with Pierce LTQ Velos ESI Positive Ion Calibration Solution (Thermo Fisher Scientific, Waltham, MA). The MS was operated using the following parameters: resolution, 100,000; isolation width (MS/MS), 1 *m/z;* normalized collision energy (MS/MS), 35; activation q value (MS/MS), 0.4; activation time (MS/MS), 30 ms. Data analysis was conducted using the Qualbrowser application of Xcalibur software (Thermo Fisher Scientific, Waltham, MA). HPLC-grade reagents were used to prepare samples for mass spectrometric analyses.

## Funding Information

The authors acknowledge the Division of Chemical Sciences, Geosciences, and Biosciences, Office of Basic Energy Sciences of the U.S. Department of Energy through Grant DE-FG02-02ER15296 (to W.W.M) for the genetic and physiological studies in *Methanosarcina acetivorans;* the National Institutes of Health grant GM097142 (to D.A.M) for mass spectrometry experiments; and the Carl R. Woese Institute for Genomic Biology postdoctoral fellowship (to D.D.N). D.D.N is currently a Simons Foundation fellow of the Life Sciences Research Foundation. The funders had no role in study design, data collection and interpretation, or the decision to submit the work for publication.

## Acknowledgements

We thank Graham A. Hudson (D.A.M. lab) for technical assistance with the HR and tandem MS data acquisition.

## Supplementary Files

**Supplementary File 1**– Lists of plasmids

**Supplementary File 2** – List of target sequences for Cas9-mediated genome editing

**Supplementary File 3** – List of strains

---

**Figure 2 - Supplementary Figure 1:**
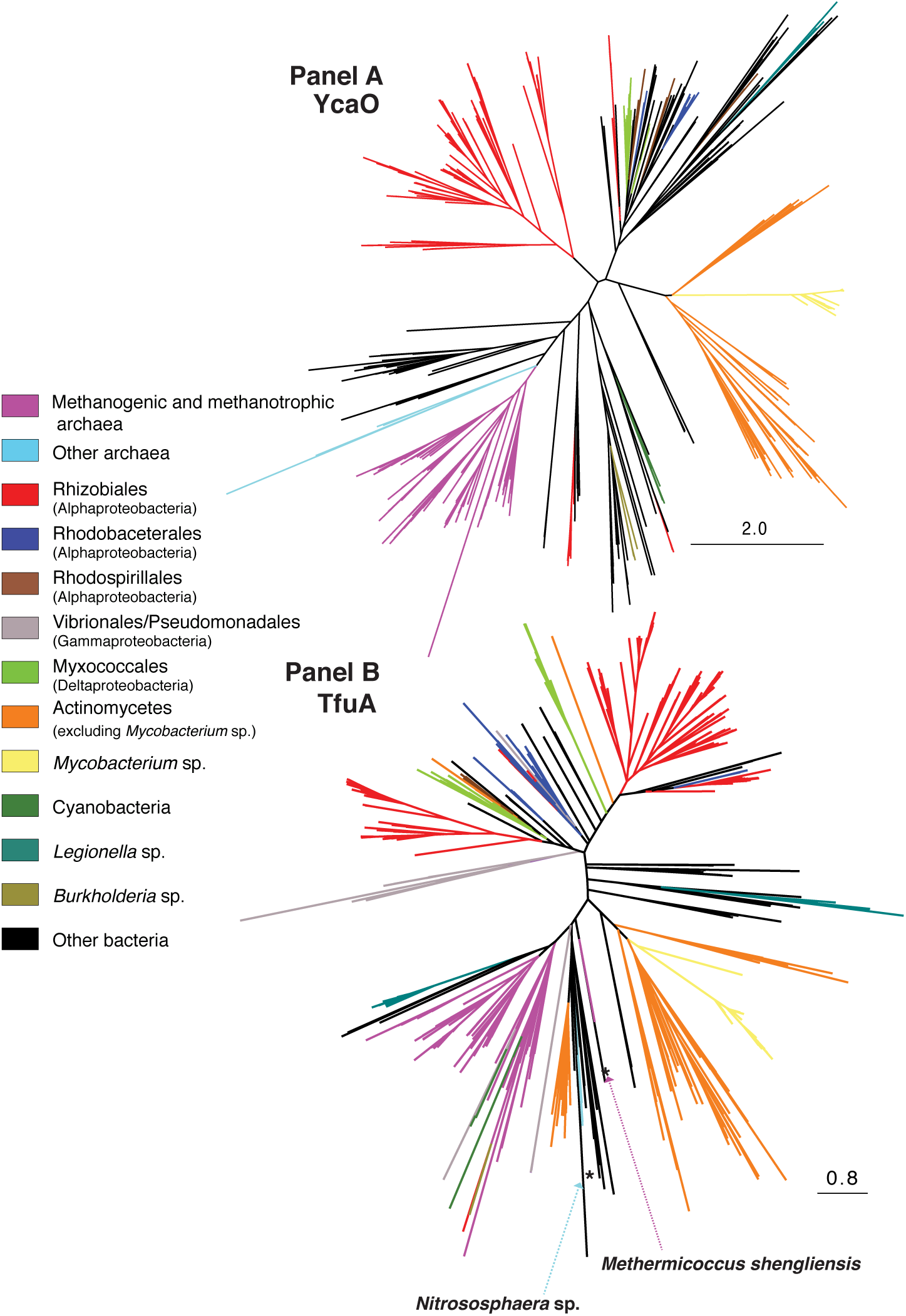
*Panel A*), Unrooted maximum-likelihood phylogeny of 1,000 YcaO sequences retrieved from the NCBI non-redundant protein sequence database using the corresponding sequence from *Methanosarcina acetivorans* as a search query. **Panel B,** Unrooted maximum-likelihood phylogeny of 1000 TfuA sequences retrieved from the NCBI non-redundant protein sequence database using the corresponding sequence from *M. acetivorans* as a search query. Divergent archaea-derived sequences on the TfuA tree are denoted with an *

**Figure 2 - Supplementary Figure 2:**
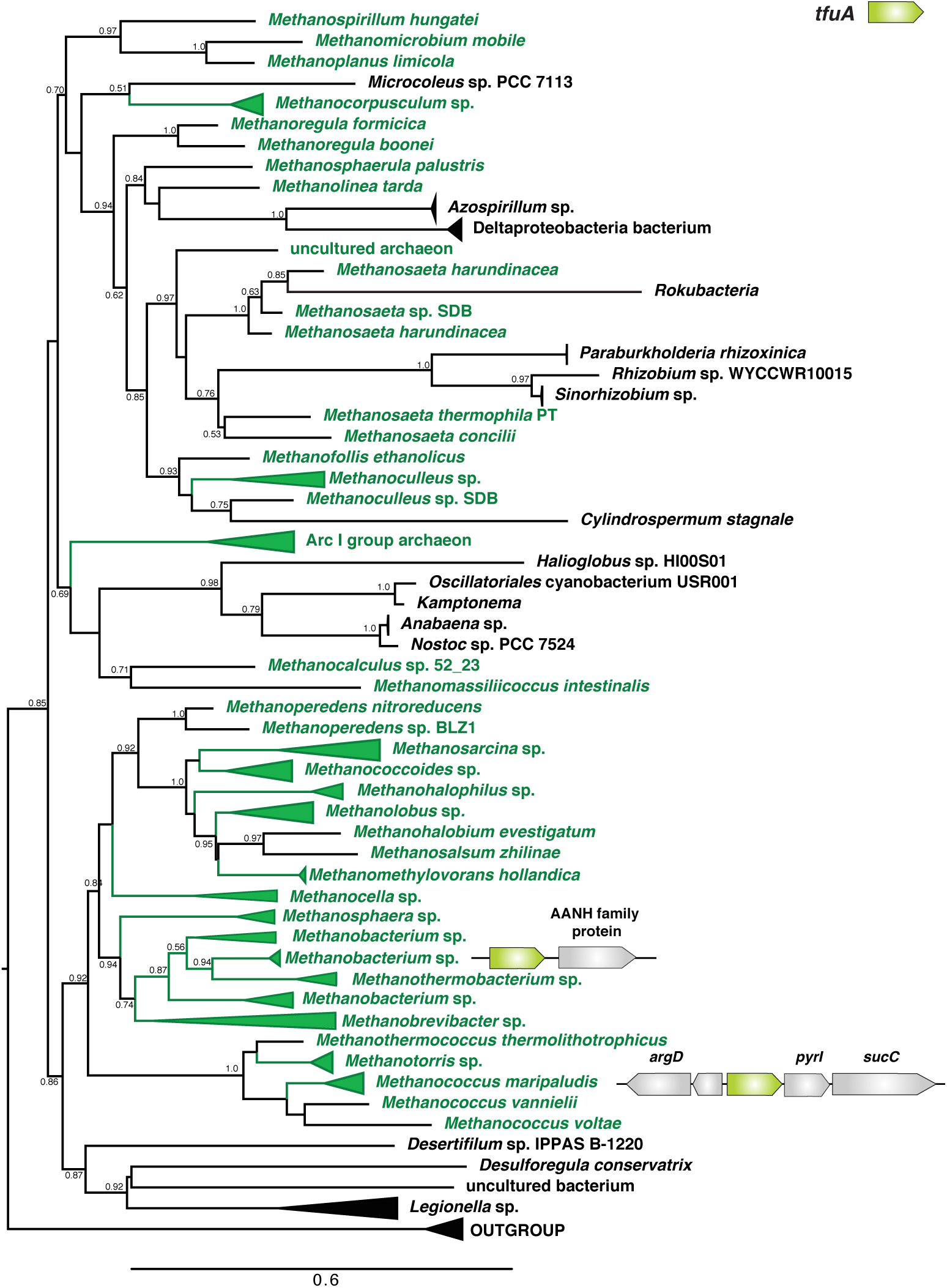
Unrooted maximum-likelihood phylogeny of 1,000 TfuA sequences retrieved from the NCBI non-redundant protein sequence database using the corresponding sequence from *Methanosarcina acetivorans* as a search query. For taxa where *tfuA* and *ycaO* are not adjacently encoded genes a gene architecture diagram is shown. The node labels indicate support values calculated using the Shiomdaira-Hasegawa test using 1000 resamples. Support values less than 0.5 have not been shown. The outgroup is comprised of various bacterial TfuA protein sequences.

**Figure 3 - Supplementary Figure 1:**
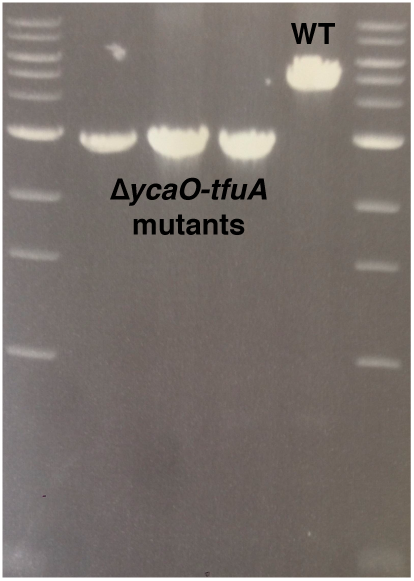
A PCR-based screen to genotype the *ycaO-tfuA* locus in *Methanosarcina acetivorans*. The PCR amplicon from three independently derived mutants containing an unmarked in-frame deletion of *ycaO-tfuA* (Lanes 2-4) is 2980 bp whereas the corresponding amplicon from a strain containing the wild-type locus is 4742 bp (Lane 5). Lanes 1 and 6 contain a 1 kb DNA ladder.

**Figure 4 - Supplementary Figure 1:**
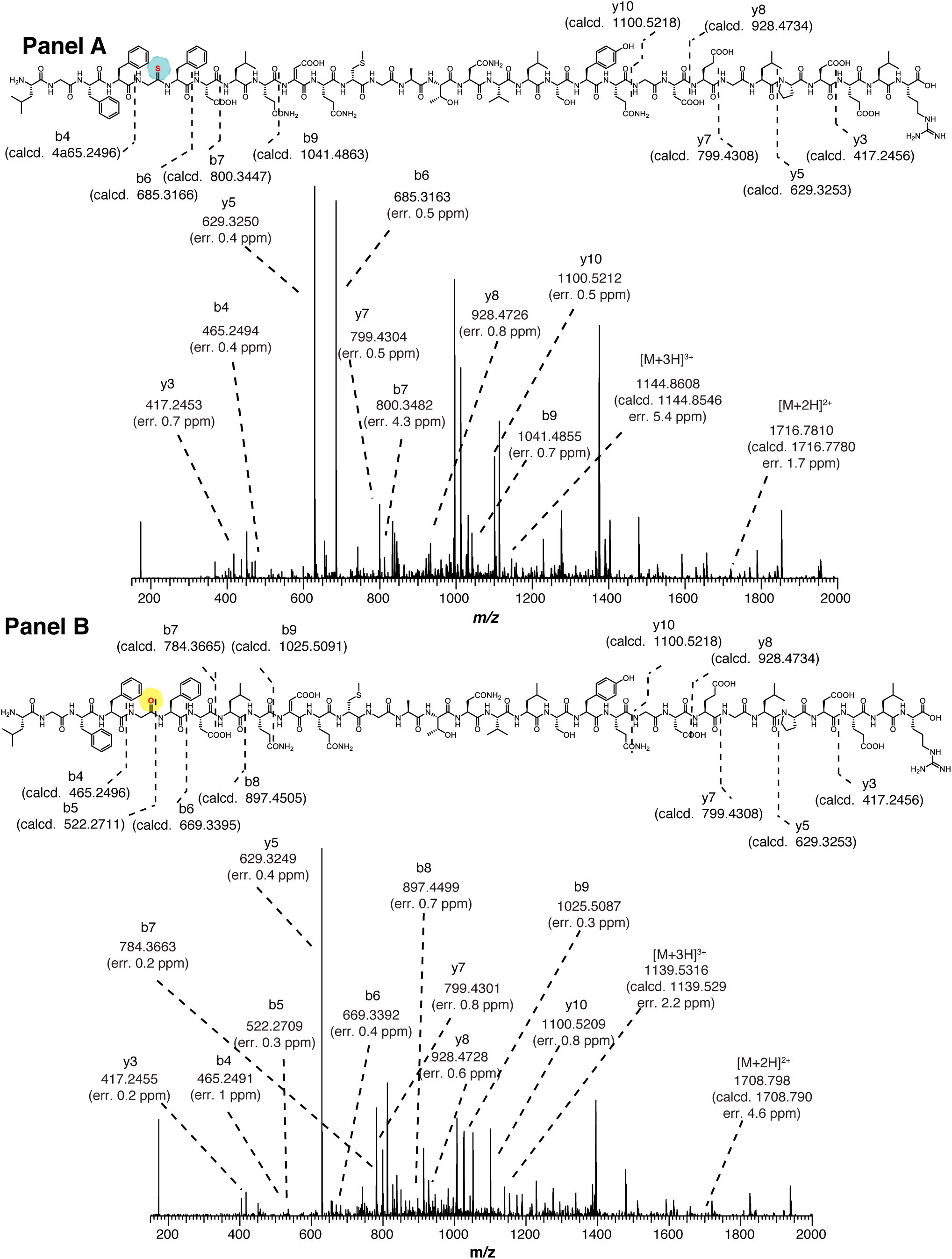
**Panel A**) High-resolutions electrospray ionization tandem mass spectrometry (HR-ESI MS/MS) analysis of a wild-type tryptic peptide from *Methanosarcina acetivorans* McrA (L461-R491, m/z 3432 Da). The b6 ion localizes the thioamide modification to Gly465. No b5 ion was detected suggesting negligible fragmentation at the thioamide bond. Predicted masses and associated errors are shown. **Panel B,** HR-ESI-MS/MS analysis of the corresponding McrA tryptic peptide from the *ΔycaO-tfuA* deletion strain of *M. acetivorans* (m/z 3416 Da). The b5 and b6 ion localizes the amide functionality at Gly465. Predicted masses and associated errors are shown.

**Figure 4 - Supplementary Figure 2:**
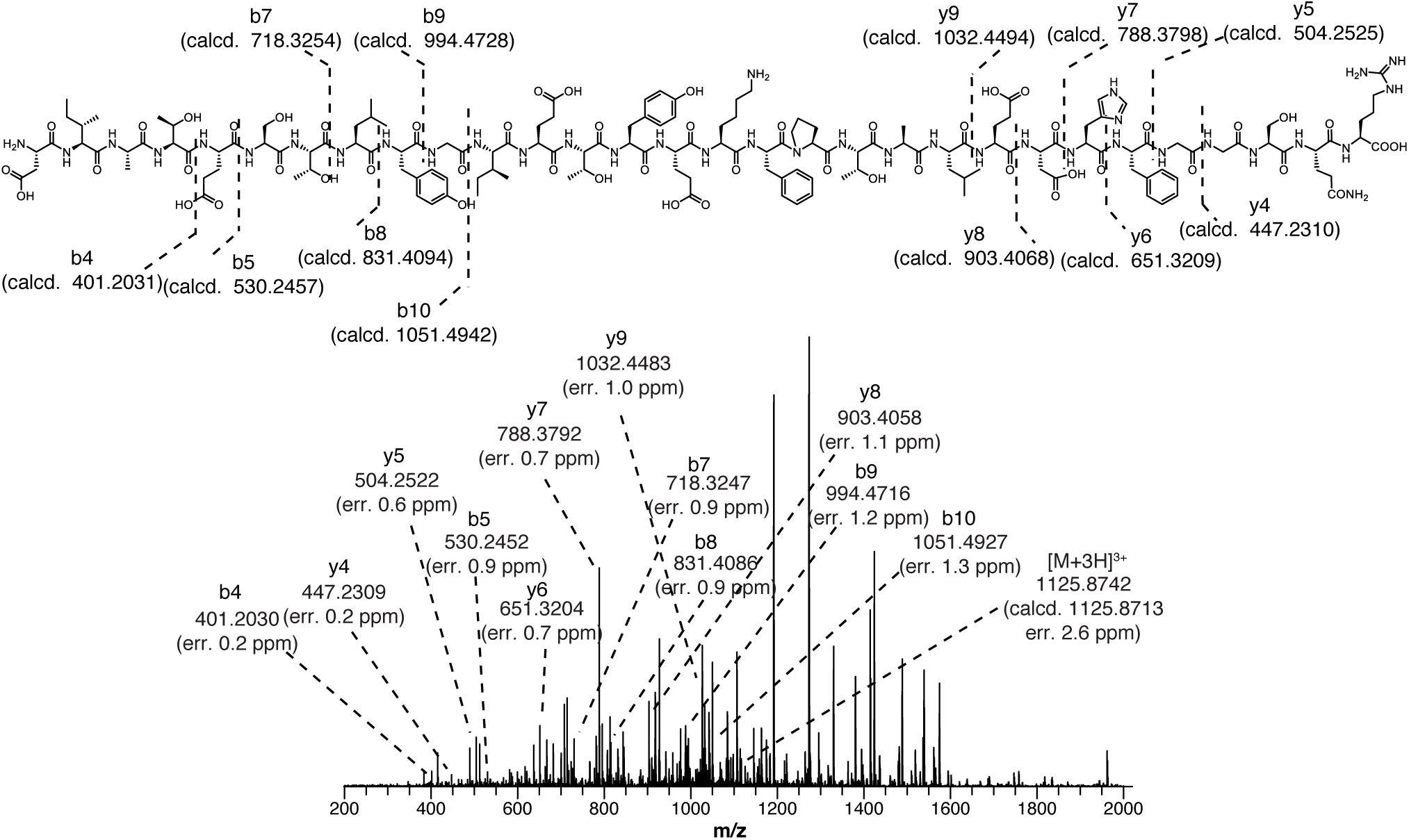
HR-ESI-MS/MS of the tryptic peptide from *Methanosarcina acetivorans* McrA (D392-R421, m/z 3375 Da) present in both the wild-type and *ΔycaO-tfuA* deletion strain. Predicted masses and associated errors are shown.

**Figure 4 - Supplementary Figure 3:**
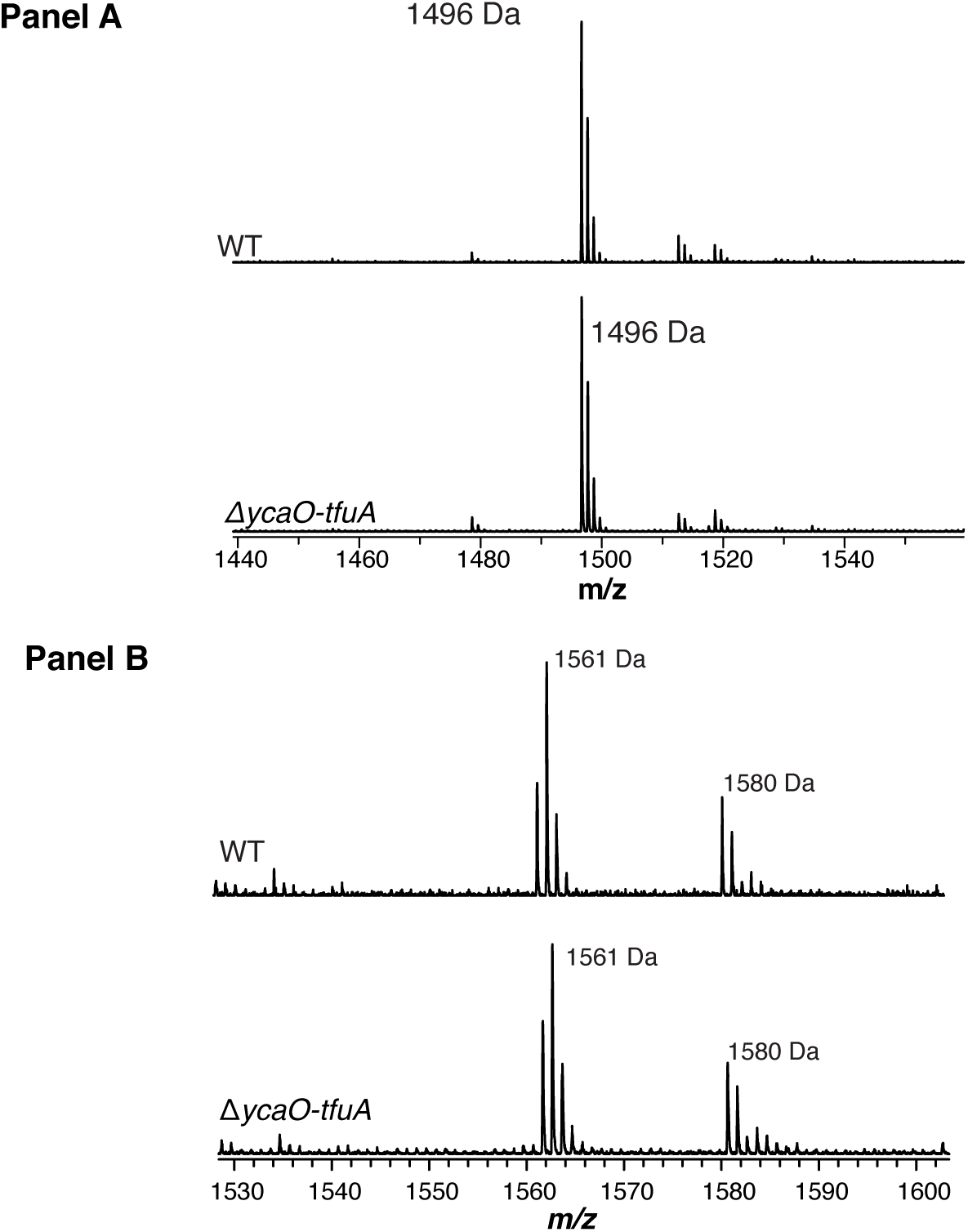
**Panel A,** Matrix-assisted laser desorption/ionization time-of-flight mass spectrometry (MALDI-TOF MS) analysis of the wild-type (WT, top) and Δ*ycaO-tfuA* deletion strain (bottom) spectra identify the McrA tryptic peptide, H271-R284 (m/z 1496 Da), containing the known *N*-methylhistidine modification. This suggests that the thioglycine formation is not a prerequisite for this methylation event. **Panel B,** MALDI-TOF MS analysis of the WT (top) and Δ*ycaO-tfuA* deletion strain (bottom) spectra identify the McrA tryptic peptide, F408-R421, (m/z 1561 Da). This peptide has been reported to contain a β-methylation at Q420 in other methanogens (9), however, it remains unmodified in *M. acetivorans*.

**Figure 5 - Supplementary Figure 1:**
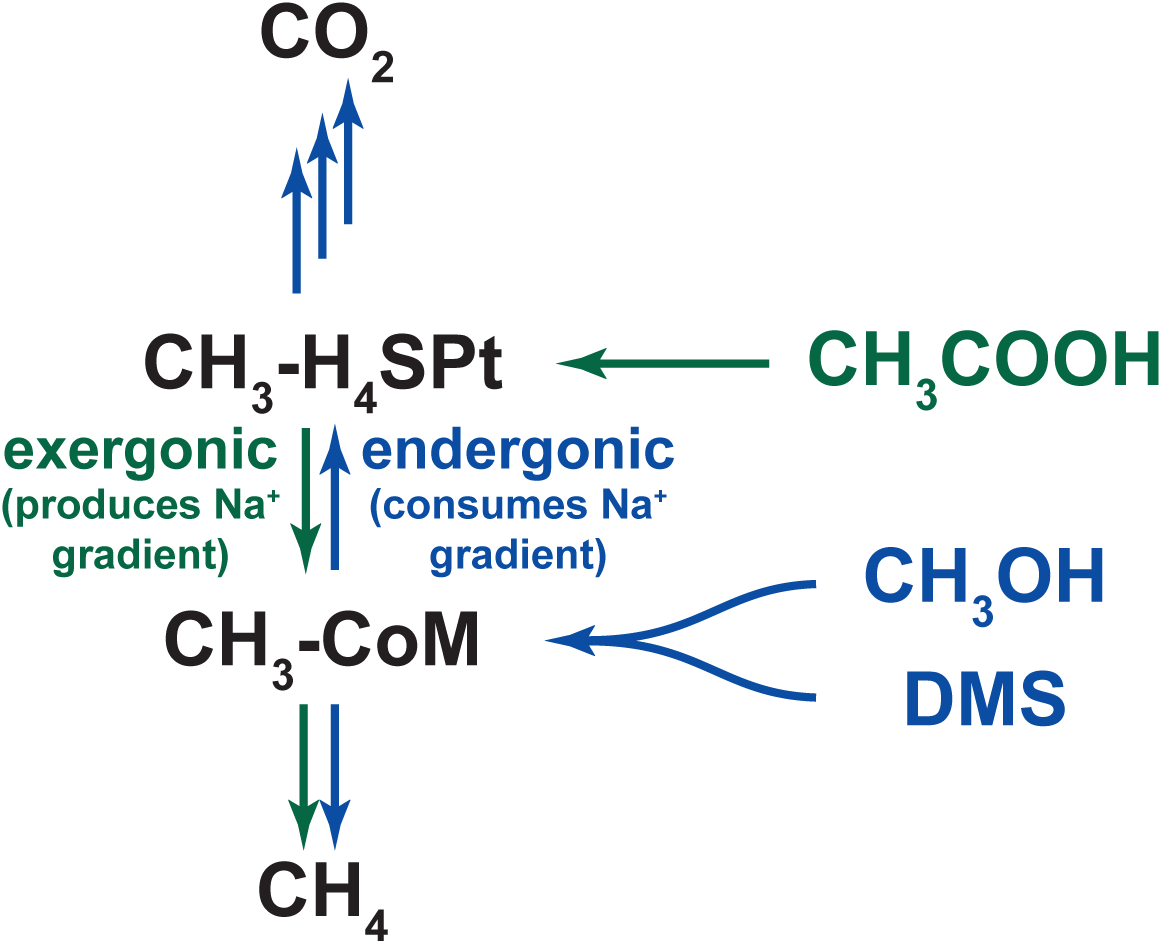
An overview of methanogenic metabolism in *M. acetivorans*. Methylotrophic substrates such as methanol (CH_3_OH) or dimethylsulfide (DMS) enter the methanogenic pathway via S-methylation of coenzyme M (CoM) and subsequent disproportionation to methane (CH_4_) and carbon dioxide (CO_2_; metabolic flux is shown as blue arrows). Notably, the first step in oxidation of CH_3_-CoM to CO_2_ is the energy-requiring transfer of the methyl moiety to generate methyl-tetrahydrosarcinapterin (CH_3_-H_4_SPt). In contrast, acetic acid enters the pathway at the CH_3_-H_4_SPt level, followed by reduction to CH_4_ (green arrows). Thus, the second step of the pathway is exergonic.

